# Learning better by learning together: dyadic visual perceptual learning on orientation discrimination

**DOI:** 10.1101/2022.06.10.495635

**Authors:** Yifei Zhang, Jian Li, Yizhou Wang, Fang Fang

## Abstract

The belief that learning can be modulated by social context is mainly supported by high-level value-based learning studies. However, whether social context can even modulate low-level learning such as visual perceptual learning (VPL) is still unknown. Unlike traditional VPL studies in which participants were trained singly, here we developed a novel dyadic VPL paradigm in which paired participants were trained with the same orientation discrimination task and they could monitor each other’s performance. We found that the social context (i.e., dyadic training) led to a greater behavioral performance improvement and a faster learning speed, compared with the single training. Interestingly, the facilitating effects could be modulated by the performance difference between paired participants. Functional magnetic resonance imaging (fMRI) results showed that, compared with the single training, social cognition areas including bilateral parietal cortex and dorsolateral prefrontal cortex displayed a different spatial activity pattern and enhanced functional connectivities to early visual cortex during the dyadic training. Furthermore, the dyadic training resulted in more refined orientation representation in primary visual cortex (V1), which was closely associated with the greater behavioral performance improvement. Taken together, we demonstrate that the social context, learning with a partner, can remarkably augment the plasticity of low-level visual information process by means of reshaping the neural activities in early visual cortex and social cognition areas, as well as their functional interplays.

## Introduction

In our daily life, people usually learn together. Learning can be facilitated in certain social contexts. For example, according to the social learning theory proposed by Albert Bandura^1^, individuals can observe others’ behaviors and learn from the feedback given to them. In addition, one’s high motivation for performing better in front of others will also facilitate learning, which is referred to as social facilitation^2–6^. It has been revealed that social cognition related areas, including dorsolateral prefrontal cortex (dlPFC), ventromedial prefrontal cortex (vmPFC), and anterior cingulate cortex (ACC), play an important role in social facilitation to learning^4,7–9^. For instance, dlPFC has been suggested to be engaged in monitoring social partners’ performance and integrating this information with one’s subjective goal during social learning. However, previous studies mainly focused on social facilitation to high-level learning processes such as value-based learning. To the best of our knowledge, whether and how social context can even affect low-level learning processes such as perceptual learning has not been studied yet.

Visual perceptual learning (VPL) is defined as a long-term improvement in visual task performance after training^10–12^. VPL is a commonly used paradigm for studying neural plasticity in human adults, and is also an effective rehabilitation therapy for visual impairment, such as amblyopia^13^, macular degeneration^14^ and cortical blindness^15^. Many studies attempted to uncover the neural mechanisms of VPL.

Electrophysiological and brain imaging studies showed that VPL could modulate neural responses in visual areas that are functionally specialized for the trained stimuli ^16–22^, resulting in greater response amplitude^18,23^, shorter response latency^24,25^, sharpened tuning function^16,20,26^, stronger external noise filtering, and internal noise suppression^27–30^. Recent studies also suggest that VPL could be mediated by higher cortical areas, which is directly involved in decision-making processes, such as the lateral intraparietal sulcus^31,32^, ACC^33^, and dlPFC. Reweighted or optimized connections between these decision-making areas and visual areas were found to be associated with better visual task performance after training ^32,34^. However, whether social cognition related areas can be engaged in VPL if subjects are trained in a social context still remains unknown.

To address the questions, we developed a novel dyadic VPL paradigm, where paired partners are trained together on a basic visual task – an orientation discrimination task, and monitor each other’s performance. By combining psychophysics and functional magnetic resonance imaging (fMRI), we investigated whether the presence of a learning partner can facilitate VPL, including the behavioral performance improvement and the learning speed, as well as the differences in neural mechanisms between the dyadic and single VPL. In Experiment 1, psychophysical results showed that training with a partner led to greater behavioral performance improvement and a faster learning speed than training singly. In Experiment 2, we found that one’s learning could be promoted even more when his/her learning partner performed better. In Experiment 3, fMRI results demonstrate that, compared with the single VPL, the dyadic VPL was more effective in refining the orientation representation in early visual cortex. Interestingly, higher areas, including bilateral parietal cortex and left dlPFC, exhibited different spatial activation patterns during the dyadic and the single training. Stronger functional connectivities between these areas and early visual cortex were also observed during the dyadic training compared with the single training. These findings demonstrate that the social context generated from learning together can significantly boost the low-level VPL, which is implemented by the synergistic mechanisms between early visual cortex and social cognition related areas.

## Results

### Dyadic vs. single visual perceptual learning

In Experiment 1, we developed a novel dyadic perceptual learning paradigm to investigate whether and how perceptual learning could be influenced by a learning partner (Figure 1A). 15 pairs of participants underwent the three phases of Experiment 1: pre-training test, training, and post-training test. During the training phase, paired participants practiced an orientation discrimination task together (i.e., dyadic VPL). Sitting in different behavioral rooms, they viewed identical stimuli, made responses and communicated through connected computers. In a training trial, two ring-shaped gratings were presented successively. They were first asked to judge the orientation change from the first to the second grating (clockwise or counter-clockwise) independently and then received feedback – both participants were correct, both participants were wrong, or participants gave inconsistent responses. If their responses were not consistent, they were asked to make a second response to decide whether to change his/her initial choice based on his/her confidence in himself/herself and his/her partner. After the second response was made, feedback was given to each participant. The feedback could be both correct, both wrong, or which participant was correct (if the second responses were still inconsistent) (Figure 1B). In addition to the 15 pairs of participants (i.e., the dyadic training group), for comparison purpose, another 15 participants (i.e., the single training group) were recruited and trained on the same orientation discrimination task singly (i.e., single VPL). During the training phase, participants in the single training group were asked to make only one response in a trial and then received correct/wrong feedback (Figure 1C).

**Figure 1.**
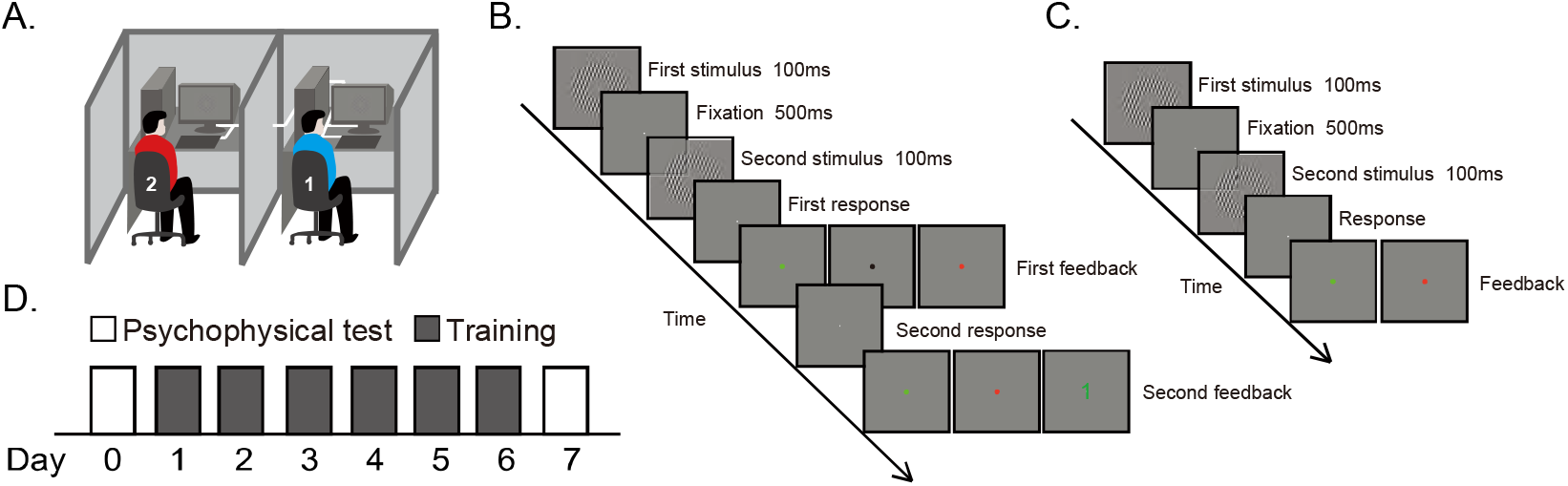
Setup, procedures, and protocol in Experiments 1 and 2. (A) Paired participants completed the dyadic training task in separate rooms in which the experimental setups for them were identical. They communicated through connected computers, rather than in a face-to-face way. (B) Experimental procedure for the dyadic training. Paired participants were asked to judge the orientation change from the first to the second grating. After they made their first and independent responses, the fixation turned green, red, or black if their first responses were both correct, both wrong or inconsistent. Paired participants were asked to make a second response if their first responses disagreed. After their second responses, the fixation turned green or red if their second responses were both correct or both wrong. If their second responses still disagreed, the ID (1 or 2) of the correct participant would be presented. (C) Experimental procedure for the single training. Participants in the single training group made only one response and then received feedback. The fixation turned green or red if the response was correct or wrong. (D) Protocol in Experiments 1 and 2. Participants underwent six daily training sessions. Psychophysical tests were performed before and after training. In Experiment 2, high-aptitude learning partners received extra single training (4 daily sessions) before the dyadic training.

For both the dyadic and single training groups, the training phase consisted of 6 daily sessions of 1040 trials (Figure 1D). For each daily session, using the method of constant stimuli, participants’ orientation discrimination thresholds were estimated based on their independent responses after the presentation of the two gratings, which reflected their individual visual aptitude. One day before and after the training phase, pre- and post-training tests were conducted to measure participants’ individual orientation discrimination thresholds also with the method of constant stimuli. By comparing the thresholds measured in the pre- and post-training tests, we could evaluate participants’ performance improvement induced by training.

The orientation discrimination thresholds in the pre-training test were submitted to a 2 × 2 mixed-design ANOVA with a between-subject factor of group (single and dyadic) and a within-subject factor of orientation (trained and untrained). The main effects of orientation (F(1,43)=1.439, p>0.1, *η*^2^ = 0.032) and group (F(1,43)=0.034, p>0.1, *η*^2^ = 0.001) were not significant, and the interaction between group and orientation (F(1,43)=0.056, p>0.1, *η*^2^ = 0.001) was not significant either, demonstrating that the initial performances of the single and dyadic training groups were comparable.

Figure 2A shows the learning curves of the single and dyadic training groups. During the training phase, the dyadic training group exhibited better task performance and a faster learning rate, compared with the single training group. The discrimination thresholds in the dyadic training group were significantly lower than those in the single training group. This performance superiority lasted from the first to the last training sessions (all six t(43)>1.69, p<0.05, Cohen’s d>0.47, one-tailed). Though the learning curves saturated after the third training session, the curve of the dyadic training group was steeper than that of the single training group in the early training phase. The discrimination thresholds were submitted to a 2×4 mixed-design ANOVA with a between-subject factor of group (single and dyadic) and a within-subject factor of day (pre-training test and training sessions 1-3). The interaction between group and day was significant (F(3,129)=2.714, p=0.048, *η*^2^ = 0.059), indicating that the dyadic training group learned faster than the single training group in the early training phase.

**Figure 2.**
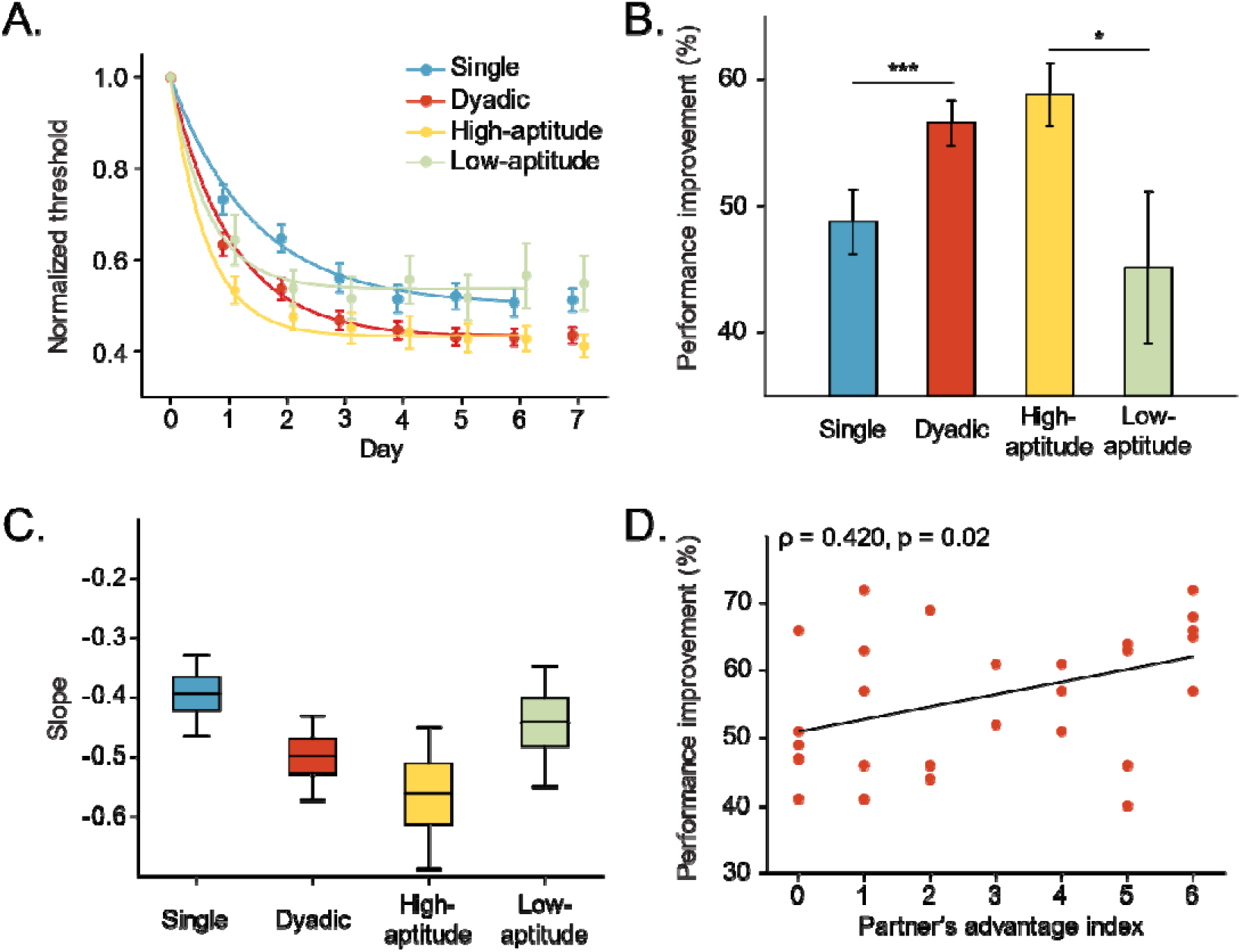
Results of Experiments 1 and 2. (A) Learning curves of the single training group, the dyadic training group, the high-aptitude group, and the low-aptitude group in Experiments 1 and 2. Discrimination thresholds are plotted as a function of training day. (B) Performance improvements in orientation discrimination threshold for the four training groups (*p<0.05, ***p<0.001). (C) Boxplots of the estimated slope for the four training groups generated by bootstrap-resampling for 5000 times. Box-whiskers are drawn down to the 5th percentile and up to the 95th percentile. (D) Correlation between participants’ performance improvement and their learning partner’s advantage index. Error bars denote 1 SEM across participants.

To further quantify the learning rate, the discrimination thresholds in the pre-training test and training sessions 1-3 were regressed linearly against the day. The slope of the regression line was used to quantify the learning rate. A bootstrap resampling method was used to estimate the statistical characteristics of the slope of the single and dyadic training groups. The slope of the dyadic training group was steeper (dyadic: *M*=-0.50, *SD*=0.04, 95%CI: [-0.59, -0.42]; single: *M*=-0.39, *SD*=0.04, *95%CI*: [-0.48, -0.32]; *d*=2.75, Figure 2C). A non-parameter jackknife permutation test was employed to examine the slope difference^35^, showing that the learning of the dyadic training group was faster (*p*<0.001).

In the post-training test, although tested singly, participants in the dyadic training group exhibited greater performance improvement than those in the single training group (t(43)=2.487, p=0.016, d=0.79, Figure 2B). In addition, there was no significant threshold difference between the post-training test and the last training day in the dyadic training group (t(29)=-0.51, p>0.1), indicating that the dyadic training effect could be fully preserved even when participants were tested singly.

### Impact of learning partner’s performance on dyadic VPL

Next, we investigated how a participant’s VPL (i.e., learning rate and performance improvement) was affected by his/her learning partner’s performance or visual aptitude during training. First, we found that the performance difference between paired participants during training was associated with their performance improvement induced by training. To quantify the performance difference between paired participants across the six daily training sessions, a participant’s advantage index was defined as the number of days on which his/her discrimination threshold was lower than his/her partner’s. For example, if a participant’s advantage index is 5, it means that if we compare the thresholds from the paired participants on each training day, this participant’s threshold is lower than his/her partner’s threshold on 5 of the 6 training days. Because a lower threshold reflects a better performance, the larger the advantage index, the better performance this participant would have than his/her partner during the dyadic training. We observed a significant positive correlation between participants’ performance improvement and their partner’s advantage index (spearman *ρ*=0.420, *p*=0.02, Figure 2D), suggesting that a learning partner with better performance can help the participant to improve more in the dyadic VPL.

To further evaluate the relationship between the dyadic training effects and the performance difference between paired participants, in Experiment 2, we sought to study the dyadic VPL of participants of interest who were paired with a high- or low-aptitude partner. Specifically, participants of interest (n=30) underwent the same training as that in Experiment 1. Half of the participants of interest were paired with a partner who underwent extra single training (4 daily sessions) before the dyadic training (dyadic VPL group with a high-aptitude partner, the high-aptitude group). The other half were paired with a partner who was presented with the grating stimuli embedded in white noise (dyadic VPL group with a low-aptitude partner, the low-aptitude group). These manipulations rendered the high- and low-aptitude partners perform better or worse than those participants of interest, respectively. It should be noted that, in both groups, all participants of interest and their partners did not know these manipulations. Participants of interest believed that his/her partner underwent the same training protocol as he/she did and his/her partner’s performance was due to his/her own aptitude.

Before analyzing the learning data, we first examined the validity of our manipulations. For the high-aptitude partners, 4-day single training significantly lowered their orientation discrimination thresholds (t(28)=-7.88, p<0.001, d=2.88). During the following dyadic training, the average advantage index of the high-aptitude partners was 4.67 (*SD*=1.88), showing that they had better task performance on most of the training days. For the low-aptitude partners, adding noise effectively elevated their discrimination thresholds and helped simulate a partner of low aptitude. The low-aptitude partners’ thresholds estimated in the first dyadic training session were significantly higher than their (t(14)=3.49, p=0.02, d=0.90) and their partner’s thresholds (t(28)=3.10, p=0.004, d=1.13) in the pre-training test. The advantage index of the low-aptitude partners was 0 for all pairs. Clearly, these results confirmed the validity of our manipulations.

Then we analyzed behavioral data of the participants of interest in the high- and low-aptitude groups. Figure 2A shows the learning curves of the two groups, as well as the learning curves of the single training group and the dyadic training group in Experiment 1. In the pre-training test, thresholds did not differ significantly between the high- and low-aptitude groups (F(1,28)=0.885, p>0.1, *η*^2^ = 0.031). In the early training phase, the slopes of the high- and low-aptitude groups were also estimated as in Experiment 1, which were both steeper than that of the single training group (jackknife permutation tests, both *p*<0.001), indicating initial faster learning rates of the dyadic training groups regardless partners’ performance. Compared with the slope of the dyadic training group, the slopes of the high- and low-aptitude groups were significantly steeper and flatter, respectively (jackknife permutation tests, both *p*<0.001), demonstrating the fastest learning rate of the high-aptitude group. As the training proceeded, the high-aptitude group rather than the low-aptitude group, showed superiority over the single training group. In the post-training test, the performance improvement of the high-aptitude group was significantly greater than that of the low-aptitude group (t(28)=2.1, p=0.045, d=0.77, Figure 2B). But there was no significant difference between the single training group and the low-aptitude group (t(28)=-0.618, p>0.1) as well as between the dyadic training and the high-aptitude group (t(43)=0.72, p>0.1). Taken together, these results demonstrate that one’s learning rate and performance improvement can be impacted by his/her learning partner’s performance.

### Neural substrates of enhanced learning effects in dyadic VPL

In Experiment 3, another 30 participants were recruited to investigate the neural substrates of the enhanced learning effects in the dyadic VPL. We sought to address two questions. One was the neural underpinnings of the enhanced performance improvement of the dyadic VPL in the post-training test, the other was the differences in neural activity and functional connectivity between the dyadic and single VPL during training.

To this end, participants were randomly split into two groups: the single training group (n=10) and the dyadic training group (n=20, 10 pairs of participants). All participants underwent three phases: pre-training test, training, and post-training test. During each test phase, a psychophysical test was first conducted to measure the orientation discrimination thresholds at 0°, 45°, and 90° deviated from the trained orientation all either clockwise or counter-clockwise (hereafter referred to as 0°, 45°, and 90°). The psychophysical test was identical to that in Experiment 1. One day after the psychophysical test, all participants in the single training group and one of each pair of participants in the dyadic training group (10 participants in each group) underwent an fMRI test (Figure 4C). They were asked to perform the orientation discrimination task at 0°, 45°, and 90° during scanning, so we could investigate how training affected the orientation representations in the brain.

The training phase consisted of five daily training sessions. For the dyadic training group, paired participants finished 5 single training runs and 5 dyadic training runs in the first daily training session, so we could make a within-subject comparison and identify the neural activity differences between the two kinds of training. In the dyadic training runs, participants of interest were scanned when they practiced the training task. His/her partner practiced this task together, but outside the scanner (Figure 3A). In the single training runs, participants of interest were scanned when they practiced the training task singly. Except the first training session, the other 4 training sessions were completed in a behavioral room. Paired participants received the dyadic training similar to that in Experiment 1. For the single training group, participants completed 10 single training runs in the scanner in the first daily session. And in the other 4 training sessions, they received the single training identical to that in Experiment 1 in a behavioral room.

**Figure 3.**
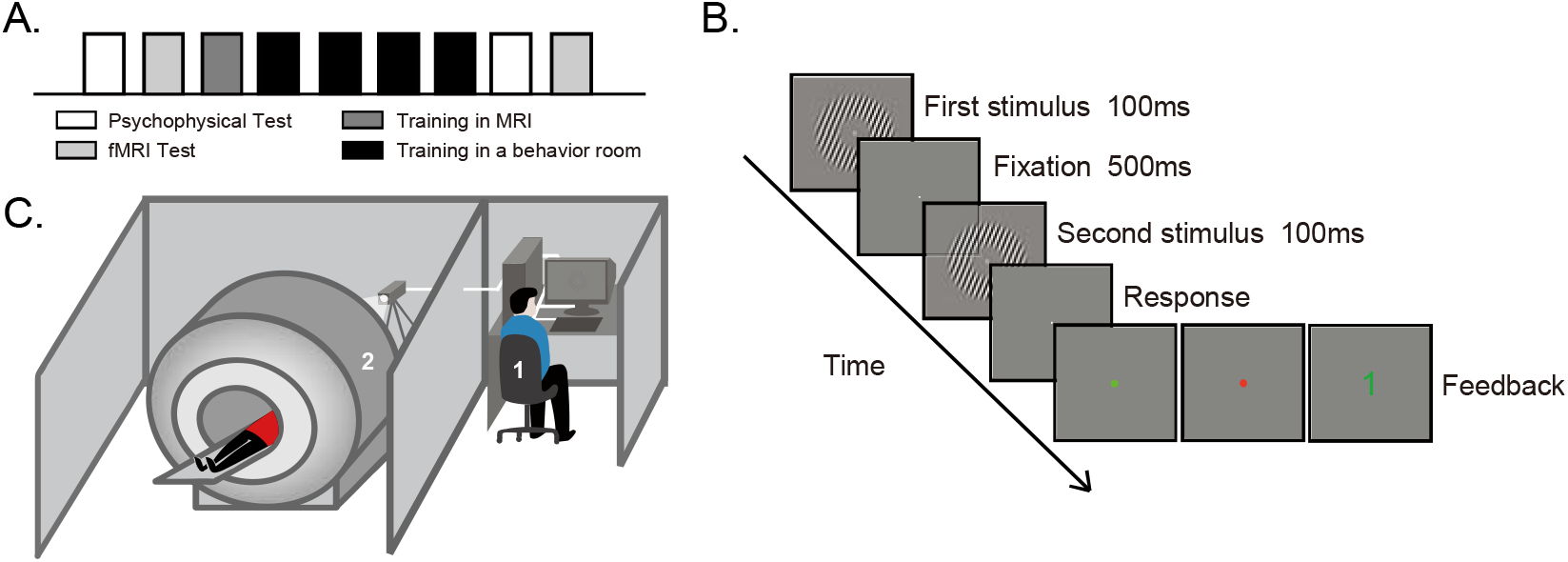
Protocol, procedure, and setup in Experiment 3. (A) Protocol. Participants underwent five daily training sessions. The first training session for participants of interest in the dyadic training group was carried out in the scanner. Psychophysical and fMRI tests were performed before and after training. (B) Experimental procedure for the dyadic training. (C) Experimental setup in the first training session. Participants of interest in the dyadic training group were scanned when he/she practiced the orientation discrimination task together with his/her partner outside the scanner, and they were also scanned when they practiced the task singly.

**Figure 4.**
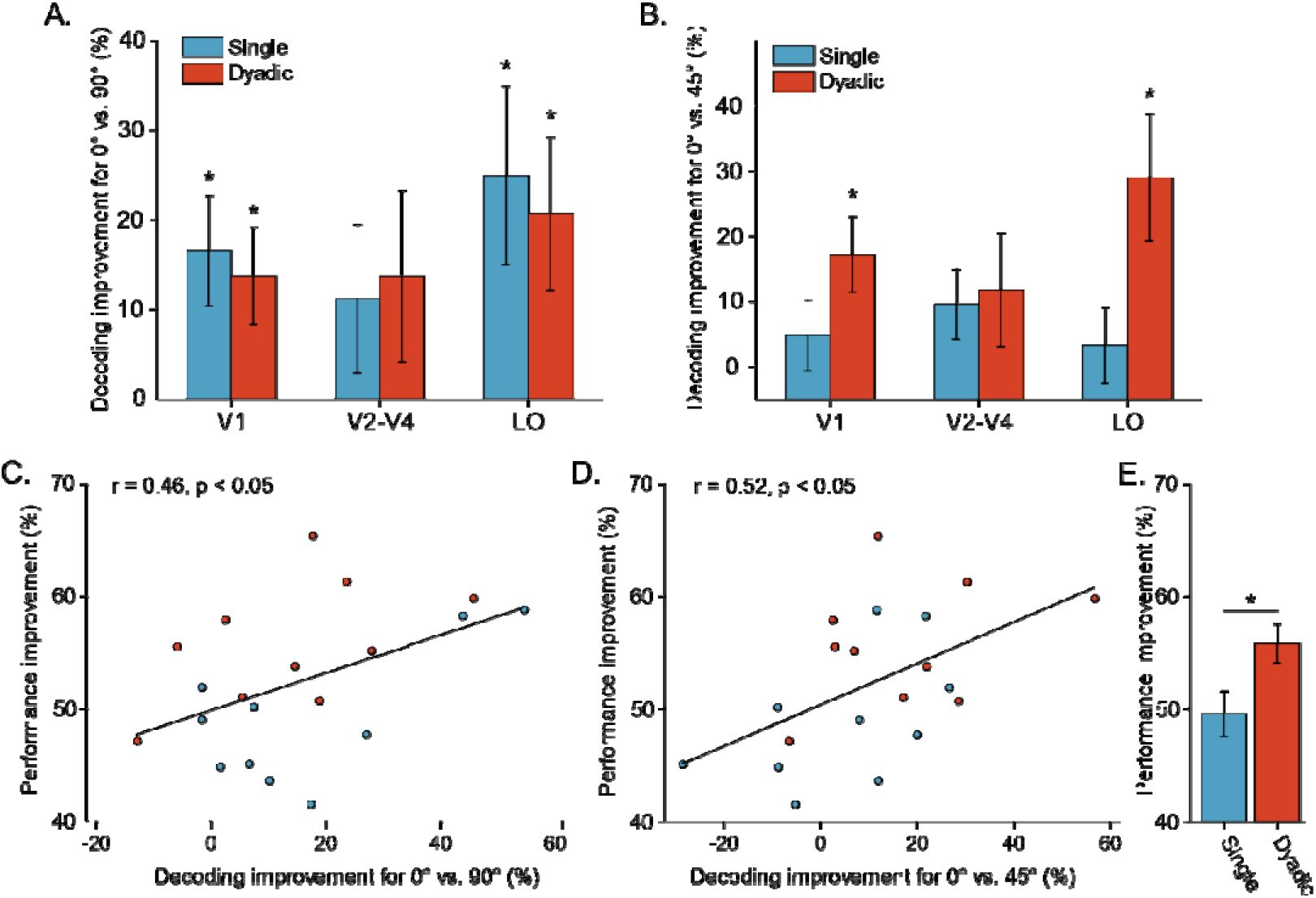
Results of Experiment 3. (A and B) Performance improvements for decoding 0° vs. 90° (A) and 0° vs. 45° (B) in visual areas (*p<0.05). (C and D) Correlations between behavioral performance improvement and decoding performance improvement in V1 for 0° vs. 90° (C) and 0° vs. 45° (D). (E) Behavioral performance improvements after the dyadic and single training (*p<0.05). Error bars denote 1 SEM across participants.

To better compare the single training and the dyadic training, in this experiment, we simplified the dyadic VPL paradigm in Experiment 1. We removed the second response and gave paired participants the final feedback after their first and independent responses. They could still know each other’s performance through the feedback (Figure 3B). We found that the simplified paradigm led to a similar learning facilitation effect as the paradigm used in Experiment 1. In the post-training test, participants in the dyadic training group gained more improvement than those in the single training group (t(18)=2.63, p=0.02, d=1.17, Figure 4E).

We used the MVPA decoding analysis to investigate the orientation representation in each retinotopic visual area. A function localizer was used to define the regions of interest (ROIs) that corresponded to the retinotopic representation of the ring-shaped gratings in V1, V2, V3, V4, and LO^36–38^. Then, we calculated the decoding accuracies for 0° versus 90° in the pre- and post-training tests. We found that, in both groups, the decoding accuracy improvements after training were significant in V1 (dyadic: t(9)=2.53, p=0.03, d=0.80, single: t(9)=2.69, p=0.02, d=0.85) and LO (dyadic: t(9)=2.43, p=0.04, d=0.76, single: t(9)=2.51, p=0.03, d=0.79, Figure 4A), but not in V2-V4 (dyadic: t(9)=1.44, p=0.19, d=0.45, single: t(9)=1.35, p=0.21, d=0.42). Since there was no significant difference in decoding accuracy among V2, V3, and V4, we pooled the accuracies in these three areas together. Notably, the improvement of V1 decoding accuracy was correlated with the improvement in behavioral performance (r=0.46, p=0.04, Figure 4B). These results are in line with previous findings that perceptual learning gave rise to more refined responses to the trained orientation in the visual cortex^20^. However, the decoding analysis for 0° versus 90° above did not reveal any difference between the single and the dyadic VPL. Therefore, the decoding analysis for 0° versus 45° was performed. Relative to 0° and 90°, the neural representations of 0° and 45° are supposed to be more similar, so it is more difficult to decode these two representations. We found that significant decoding improvement only occurred in V1 (t(9)=2.99, p=0.02, d=0.94) and LO (t(9)=2.99, p=0.02, d=0.94, Figure 4C) in the dyadic training group. No significant improvement was found in the single training group (all *p*>0.1). The improvement of V1 decoding accuracy can also predict the behavioral performance improvement (r=0.52, p=0.02, Figure 4D). These findings suggest that, compared with the single training, the dyadic training could further refine the neural representation of the trained orientation, which might explain the behavioral enhancement in the dyadic training group.

Next, we investigated the differences in neural activity and functional connectivity between the single and the dyadic training runs. Univariate GLM analysis did not yield any cluster, showing different overall BOLD activities between the dyadic and single training. Then, searchlight analysis was used to uncover a cluster or clusters, showing different spatial activation patterns between these two kinds of training. This analysis identified bilateral Intraparietal lobe (IPL) and left dorsal lateral prefrontal cortex (Figure 5A, rIPL: 40,-62,44; lIPL: -40, -58, 44; ldlPFC: -36, 54, 0). These areas were further set as seeds to investigate the functional connectivity to other brain areas by using the psychophysiological interaction (PPI) analysis method. We found that both bilateral IPL and ldlPFC showed enhanced functional connectivity to early visual cortex during the dyadic training, compared to the single training (left EVC: -24, -96, -2; right EVC: 22, -94, 2; p<0.01, uncorrected, Figure 5B-J). The left medial frontal gyrus (lMFG, -12, 0, 58) was also identified by the searchlight analysis. However, the PPI analysis seeded at lMFG did not identify a statistically significant region in visual cortex. Taken together, these analyses demonstrate that the functional couplings between early visual cortex (including V1) and higher areas (e.g., IPL and dlPFC) were enhanced during the dyadic training.

**Figure 5.**
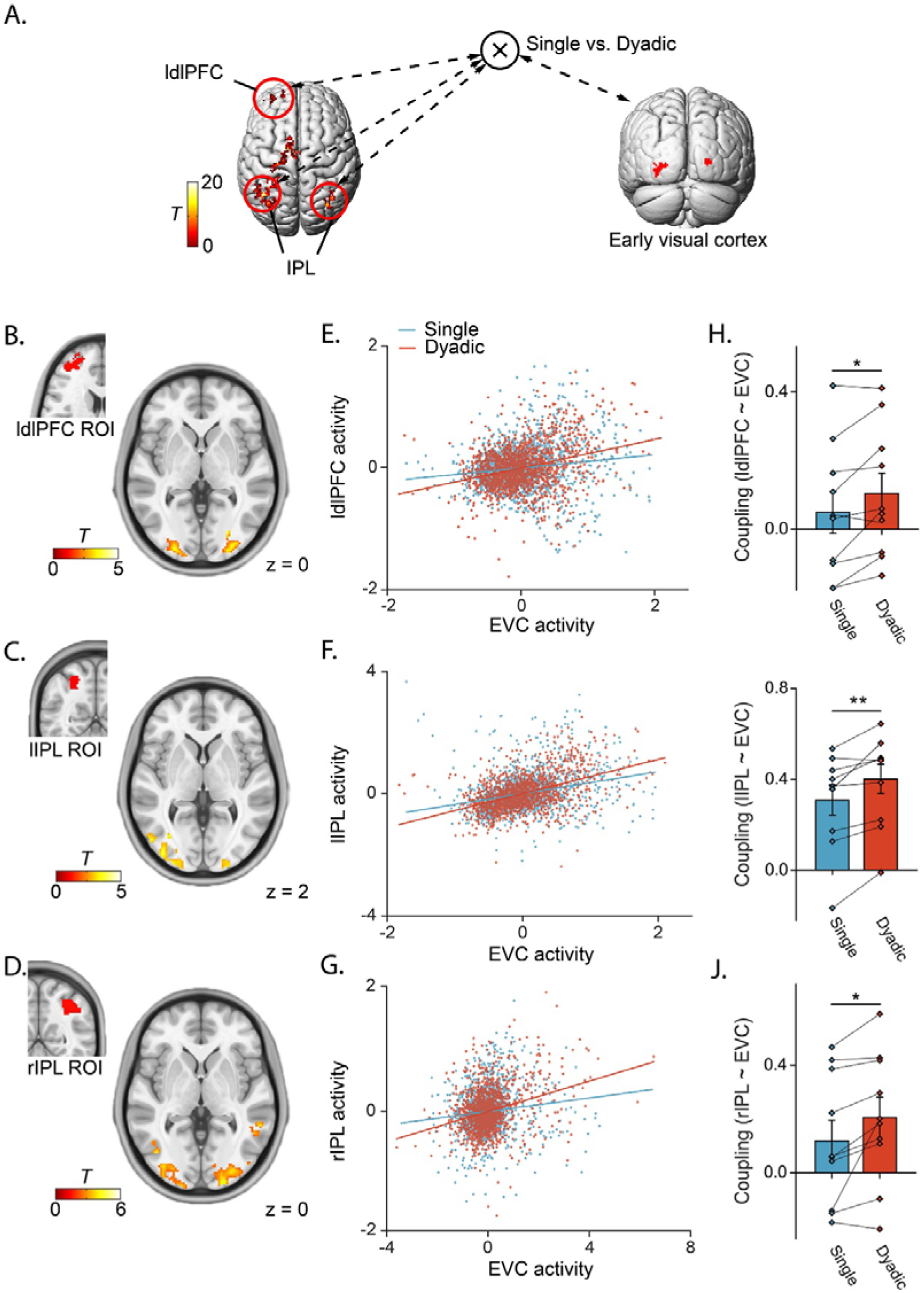
Different spatial activity patterns and functional connectivities during the dyadic and single training. (A-D) Results of searchlight and PPI analyses. Bilateral parietal cortex and left dorsolateral prefrontal cortex showed different spatial activity patterns during the dyadic and single training (A). Compared with the single training, PPI analysis identified increased functional connectivities between ldlPFC (B), lIPL (C), rIPL (D) as seed regions and early visual cortex as the target region during the dyadic training. (E-G) Correlations of activities in the target and seed regions for the dyadic and single training in a representative participant. (H-J) Boxplots of coupling strength across all participants for the dyadic and single training (*p<0.05, **p<0.01). Error bars denote 1 SEM across participants.

## Discussion

In this study, we developed a dyadic VPL paradigm to investigate VPL in the presence of a learning partner. In this social context, paired participants underwent the orientation discrimination training, during which one could monitor his/her partner’s performance at the end of each trial via connected computers. Using this paradigm, in Experiment 1, we found that learning with a partner could significantly facilitate VPL, resulting in a greater behavioral performance improvement and a faster learning speed. In Experiment 2, we found that the performance difference between paired partners could modulate the behavioral performance improvement. In Experiment 3, the fMRI results demonstrate that, relative to the single VPL, the dyadic VPL gave rise to more refined orientation representations in V1 and LO. Such refinements might be due to the interplay between early visual areas and higher areas including IPL and dlPFC, in which different spatial activation patterns and functional connectivities to early visual cortex were observed during the single and the dyadic training.

It might be argued that the greater behavioral performance improvement and the faster learning speed could be simply due to more feedbacks and responses, namely, the feedback of inconsistency and a second response, in the dyadic VPL, rather than the social context. The results obtained from Experiments 2 and 3, however, argue against this explanation. It can be seen in Experiment 2 that the performance of participants paired with a low-aptitude partner did not exhibit the VPL facilitations although they were given more feedbacks and made a second response, compared with the single training. In Experiment 3, participants in the dyadic training group still exhibited similar VPL facilitations to those in Experiment 1, even though the second response was removed, which made the numbers of feedback and response equal between the single and the dyadic VPL groups.

In the dyadic VPL, the presence of a learning partner generated a context for social facilitation^39,40^. First, compared with performing the training task singly, one would have higher motivation and alertness to perform better in front of a learning partner. Second, when inconsistent responses appeared, one would gradually realize the performance difference between him/her and his/her partner throughout the training process, which would lead to social comparison. Learning with a high-aptitude partner could result in more feelings like “My partner is correct and I am wrong”, which would threat one’s self-esteem. Such an upward comparison might give rise to higher motivation which could stimulate the participants to concentrate and improve their performance more^41,42^. On the other hand, the downward comparison when learning with a low-aptitude partner might protect one’s self-esteem and attenuate the social facilitating effect, thus leading to a comparable improvement to the single training^43^.

Converging evidence shows that motivation and social comparison are associated with several key brain areas, including dlPFC, IPL, vmPFC, dorsal ACC and striatum^39,44^. Especially, dlPFC has been considered as a pivotal hub to integrate information from different sources^45^, make strategic responses^46,47^and motivate behaviors^7,8^. For instance, McDonald et al. found that dlPFC displayed selective activation when subjects’ actions were highly opponent-sensitive in real-time video game playing^47^. A lesion study from Mah et al.^48^ also showed that dlPFC lesion could lead to impairment in social perception. Besides integrating contextual information, a study from Ballard et al. demonstrated that dlPFC was the exclusive entry point to send the signals of anticipated reward to the mesolimbic and dopamine system, thereby initiating motivated behaviors. IPL plays a critical role in the computation of social comparison, especially, the representation of the attributions of self and others^49,50^. Applying rTMS to create a ‘virtual lesion’ over right IPL could causally disrupt self-other discrimination^51^. In this study, searchlight and functional connectivity analyses identified different dlPFC/IPL spatial activation patterns and different dlPFC/IPL-EVC functional connectivities during the dyadic and the single training, which might reflect the social comparison process and the enhanced motivation-related modulation during the dyadic training.

VPL can occur at multiple cortical areas and manifest in various forms^11^. VPL associated with high-level cognitive functions such as attention and reward has been shown to manifest as connectivity reweighting between sensory and higher areas^32,33^. Our fMRI experiment demonstrate that the enhanced plasticity induced by the dyadic VPL, relative to the single VPL, could occur in early visual cortex. We first replicated a previous finding that VPL can refine the orientation representation in V1^20^. Such a representation refinement in V1 was closely associated with the behavioral performance improvement. What is new here is that the orientation representation in LO, an area associated with complex visual pattern and object processing, could also be refined by VPL. Furthermore, we found that, compared with the single training, the dyadic training was more effective in refining the orientation representations in V1 and LO, which contributed to the greater behavioral performance improvement. These results suggest that high-level social cognition processes such as social facilitation could even impinge early visual cortex and shape the characteristics of its neural responses. Our study is reminiscent of previous findings that other high-level cognitive processes, such as reinforcement, could influence neural activity in early visual cortex. Behavioral studies from Seitz et al.^52^ and Wang et al.^53^ consistently found that reward-evoked VPL could not transfer from the trained eye to the untrained eye, indicating a neural change occurring at an early stage of the visual processing hierarchy. A neuroimaging study from Arsenault et al.^54^ found that the dopaminergic reward signal could selectively shape neural activity in early visual cortex.

Finding new ways to augment brain functions effectively and rapidly is a perennial issue in medical and neuroscience communities^55^. In the past decades, numerous studies have shown that VPL can reliably enhance long-term visual abilities and have irreplaceable advantages in treating neuro-ophthalmic disorders^10,13,15,56^. To achieve greater effectiveness and better clinical applications, several add-on methods combined with visual training have been proposed to enhance VPL, including transcranial electricity stimulation (tES) and adjunctive medications. Yang et al.^57^ found that direct current stimulation (tDCS) over visual cortex could improve behavioral performance by facilitating awake consolidation of VPL. He et al. found that 10Hz transcranial alternating current stimulation (tACS)^58^ over visual cortex could help subjects acquire greater performance improvement within a shorter time. Rokem and Silver^35^ reported that the cholinesterase inhibitor donepezil could also significantly augment the magnitude of VPL. The current study introduces a new VPL paradigm and demonstrates that learning with a partner can increase the magnitude of performance improvement and accelerate the learning speed. Compared with tES and medications, this paradigm is more tolerable, less costly, and bears minimal risk of side effects, providing a possible way to treat visual and ophthalmic disorders more effectively and rapidly.

In conclusion, we found that the social facilitation resulted from learning together could remarkably boost the low-level VPL, significantly modify the spatial activity patterns in bilateral parietal cortex and left dlPFC and enhance their connectivities with early visual cortex, and profoundly refine the orientation representations in visual cortex. These findings also provide new insights for future clinical interventions and enhancements on visual and even broader cognitive functions.

## Acknowledgement

We thank Dr. Qian Wang, Xiaoqing Wang, and Dr. Tao He for their helpful comments on the draft. This work was supported by the National Science and Technology Innovation 2030 Major Program (2022ZD0204802), the National Natural Science Foundation of China (31930053) and Beijing Academy of Artificial Intelligence (BAAI).

## Methods

### Key resource table

| Reagent or resource | Source | Identifier |
| --- | --- | --- |
| Software and algorithms |  |  |
| MATLAB R2019a | The MathWorks, Natick, Massachusetts, USA | https://www.mathworks.com/ |
| Psychtoolbox 3.0.13 | <sup>59</sup> | https://www.psychtoolbox.org/ |
| SPM12 | <sup>60</sup> | https://www.fil.ion.ucl.ac.uk/spm/software/spm12/ |
| The decoding toolbox 3.997 | <sup>61</sup> | https://www.bccn-berlin.de/tdt |
| Brain Voyager QX 2.4 | Brain Innovations, Maastricht, the Netherlands | https://www.brainvoyager.com/ |
| Other |  |  |
| 3T Siemens Prisma MRI scanner | Center for MRI Research at Peking University | http://www.aais.pku.edu.cn/yjzx/ |
| 64-channel phase-array head coil | Center for MRI Research at Peking University | http://www.aais.pku.edu.cn/yjzx/ |

### Participants

There were 45 participants (20 males) in Experiment 1, 60 participants (23 males) in Experiment 2, and 30 participants (15 males) in Experiment 3. All participants were right-handed with reported normal or corrected-to-normal vision and had no known neurological or visual disorders. They were naïve to the purpose and design of this study. Their ages ranged from 18 to 27 years. They gave written, informed consent in accordance with the procedures and protocols approved by the human subject review committee of Peking University.

### Behavioral and imaging data acquisition

In the psychophysical experiments, visual stimuli were presented on a Display++ 32-inch monitor (Cambridge Research Systems Ltd; refresh rate: 100Hz; spatial resolution: 1024×768) with a gray background (30 cd/m^2^) at a viewing distance of 70 cm. A head and chin rest were used to stabilize participants’ head position.

The fMRI experiments were performed on a 3T Siemens Prisma MRI scanner at the Center for MRI Research at Peking University. MRI data were acquired with a 64-channel phase-array head coil. In the scanner, visual stimuli were back-projected via a video projector (refresh rate: 60 Hz; spatial resolution: 1024×768) onto a translucent screen placed inside the scanner bore. Participants viewed the stimuli through a mirror mounted on the head coil. The viewing distance was 70cm. Blood-oxygen-level-dependent (BOLD) signals were measured using an echo-planner imaging (EPI) sequence with a multiband acceleration factor of 2 (TE: 30ms; TR: 2000ms; flip angle: 90°; acquisition matrix size: 112×112; FOV: 224×224 mm^2^; slice thickness: 2mm; number of slices: 62; slice orientation: transversal). FMRI slices covered the whole brain. In each MRI session, a T1-weighted high-resolution 3D structural dataset were acquired for each participant before functional runs using a 3D-MPRAGE sequence (voxel size: 1×1×1mm^3^).

### Stimuli and design

Visual stimuli were ring-shaped sinusoidal gratings with their edge smoothed with a Gaussian filter (outer radius: 4°; inner radius: 1°; Michelson contrast: 1.0; spatial frequency: 2 cycles/°; mean luminance: 30 cd/m^2^). The phase of the gratings was randomized. The spatial frequency also changed slightly in each grating, which was randomly drawn from a Gaussian distribution with a mean of 2 cycles/° and a standard deviation of 0.02 cycles/°. In the dyadic training sessions in Experiment 2, the gratings presented to the ‘low-aptitude’ partner were embedded in white noise. The root-mean-square (RMS) contrast of the noise was 0.72. The RMS contrast is defined as the standard deviation of pixel luminance values divided by the mean pixel luminance^62,63^.

### Experiment 1

In Experiment 1, we investigated how dyadic training affected the learning speed and performance improvement compared with the single training. Participants were randomly assigned into the single training group (n=15) or the dyadic training group (n=30), in which they were trained on an orientation discrimination task singly or in pairs. Both groups underwent three phases: pre-training test, orientation discrimination training, and post-training test. Pre- and post-training tests took place on the day immediately before and after training. Note that all participants in Experiment 1, as well as those in Experiments 2 and 3 as seen below, were tested singly in the pre- and post-training tests.

During the training phase, each participant underwent 6 daily training sessions to perform an orientation discrimination task around a base orientation θ, which was chosen randomly as 22.5° or 112.5° (0° was the upward direction) at the beginning and was fixed for all the training sessions. Each training session consisted of 1040 trials. In a trial, two ring-shaped gratings centered at the fixation were presented for 100 ms sequentially, and the inter-stimulus interval was 500 ms. Participants were asked to indicate the orientation change from the first to the second grating (clockwise or counterclockwise). The orientation of one grating was randomly drawn from a Gaussian distribution with a mean of θ and a standard deviation of 1° to ensure that participants had to actively compare the two gratings to complete the task, rather than to rely on the remembered base orientation. The orientation change could be 0°, ± 0.5°, ±0.8°, ±1.4°, ±2.4°, ±4°, or ±8°, with 80 trials for each of the thirteen orientation changes. In the single training group, after participants made their responses, informative feedback was provided by changing the color of the fixation to green (correct response) or red (wrong response) for 1000ms. The next trial began 1000ms after feedback. In the dyadic training group, after paired participants made their first responses, the first feedback was provided by changing the color of the fixation to green (both participants were correct), red (both participants were wrong), or black (inconsistent responses from two participants). The next trial began 1000ms after the first feedback if the first responses were consistent. Otherwise, paired participants were asked to make a second response to decide whether to change the first response. After the second response, the second feedback was provided. The color of the fixation changed to green or red if the second responses from two participants were both correct or both wrong. If the second responses were still inconsistent, the ID (1 or 2) of the correct participant was presented to both participants. Note that subjects’ ID was assigned at the beginning of this experiment.

In the pre- and post-training tests, participants’ orientation discrimination thresholds were measured around the two base orientations (i.e., 22.5° and 112.5°) using the method of constant stimuli (same as above). 832 trials without feedback were completed for each base orientation, with 64 trials for each of the thirteen orientation changes.

### Experiment 2

In Experiment 2, we investigated how participants’ VPL effects were affected by the performance of their partner during training. Here, participants were randomly assigned into two dyadic training groups – the high-aptitude group and the low-aptitude group, with 30 participants in each group. One of the paired participants was the participant of interest and the other received an experimental manipulation to become a partner with high- or low-aptitude. The experimental design and the stimuli presented to the participants of interest were identical to the dyadic training group in Experiment 1. In the high-aptitude group, prior to the dyadic training, the partners were asked to complete single training for 4 days of 1040 trials. This training was identical to the single training in Experiment 1, resulting in a significant performance improvement in orientation discrimination before the dyadic training. In the low-aptitude group, the partners were presented with gratings embedded in white noise during the dyadic training. The white noise increased the task difficulty and therefore impaired the task performance. In both groups, the participants of interest were ignorant of the experimental manipulation applied to their partner. They were told that the training procedure received by their partner was identical to theirs.

### Experiment 3

In Experiment 3, we investigated the neural substrates of the enhanced learning effect in the dyadic VPL. Similar to Experiments 1 and 2, Experiment 3 also consisted of three phases: pre-training test, orientation discrimination training, and post-training test. During each test phase, we first measured the orientation discrimination thresholds at 0°, 45°, and 90° deviated from the trained orientation all either clockwise or counter-clockwise (hereafter referred to as 0°, 45°, and 90°), using the method of constant stimuli in Experiment 1. One day after acquiring the thresholds, we measured BOLD signals responding to the three orientations (i.e., 0°, 45°, and 90°) in 8 fMRI runs. Each run contained 12 stimulus blocks of 12s, four blocks for each of the three orientations. Stimulus blocks were interleaved with blank blocks of 12s.

Each stimulus block consisted of six trials. In a trial, two gratings were each presented for 100 ms. They were separated by a 500 ms blank interval and were followed by an 800 ms blank interval. Participants were asked to make a 2-AFC judgement of the second orientation relative to the first one by pressing one of two buttons. The next trial began 500ms after button press. Here, the orientation change between the two gratings was the discrimination threshold measured in the psychophysical test and made participants perform equally well (75% correct) across the three orientations.

The training phase consisted of 5 daily sessions. For the dyadic training group (n=20), in the first session, all participants completed 5 single training runs and 5 dyadic training runs, but only the participants of interest were scanned. In both kinds of training runs, there were 12 stimulus blocks of 12 s, interleaved with 12 blank blocks of 12 s. The trial structure was similar to that in the fMRI test except that informative feedback was provided for 1 s after response, making each block contain only 4 trials instead of 6 trials. In the single training runs, participants completed the task singly, and feedback was provided by changing the color of the fixation to green (correct response) or red (wrong response). In the dyadic training runs, only the participant of interest was in the scanner. His/her partner completed the task together, but outside the scanner. Feedback was provided by changing the color of the fixation to green (both correct), red (both wrong), blue (correct and inconsistent with partner), or yellow (wrong and inconsistent with partner). Participants were instructed on the meaning of the feedback before scanning. At the beginning of each run, participants were informed that this run was a single or dyadic training run. In the rest 4 training sessions, paired participants underwent the dyadic training in the behavioral room. The dyadic training was similar to that in Experiment 1, except the second response part was removed. After paired participants made their first independent responses, they received the final feedback immediately. The feedback was both correct, both wrong, or which participant was correct.

For the single training group (n=10), during the first training session, participants completed 10 single training runs in the scanner. In the rest 4 training sessions, they underwent the single training in the behavioral room. The single training was identical to that in Experiment 1.

In the retinotopic mapping session, we defined retinotopic visual areas (V1, V2, V3, V4, and LO) using a standard phase-encoded method developed by Sereno et al.^36^, Engel et al. ^37^, and Wandell et al.^38^ in which participants viewed a rotating wedge that created traveling waves of neural activity in visual cortex. We also performed a localizer run to identify voxels in the retinotopic areas responding to the trained stimuli. This block-design localizer run contained 12 stimulus blocks of 12 s, interleaved with blank blocks of 12 s. A full contrast checkerboard stimulus flickering at 7.5Hz was presented in stimulus blocks. The size and the location of the checkerboard stimulus were identical to those of the trained ring-shaped gratings.

### Quantification and statistical analysis Behavioral data analysis

Behavioral data were processed using MATLAB (MathWorks), Psychtoolbox^59^, and custom scripts. To estimate the orientation discrimination thresholds, the percentage of trials in which participants made a correct response was plotted as a psychometric function of the orientation change. We used a Weibull function to fit the psychometric values and interpolated the data to find the 75% accuracy point (i.e., the orientation discrimination threshold). The behavioral performance improvement was calculated as

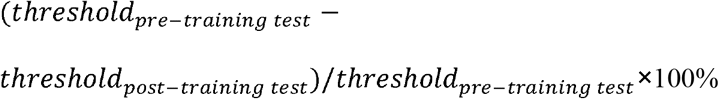

To characterize the learning process, thresholds of the pre-training test and training sessions 1-3 were regressed linearly against the day. The slope was used to quantify the learning rate. Since the data did not allow a reliable slope estimation at the single participant level, bootstrap and non-parametric permutation methods were used to examine the slope difference between groups. A bootstrap method was used to estimate the mean, variance, and confidence interval of the slope^64^. In each bootstrap sampling, we randomly sampled *n* participants with replacement (*n* was the group size), and then estimated the thresholds. After averaging the thresholds of the n sampled participants, the linear regression was performed to estimate the slope. The slope distribution was estimated by bootstrap sampling 5,000 times for each group. In the permutation test, a jackknife procedure was employed^35^. The model was fit to *n* resamples from the data. For each resample, the data from one participant were omitted, and the thresholds from the remaining *n-*1 participants were averaged. The slope was then estimated for the averaged thresholds. This produced *n* different values of slope and built a jackknife sample. The values of slope were then compared across the jackknife samples. 10,000 permuted samples were created by randomly recoding the group from which each slope was taken. The actual difference between the means of the jackknife distributions was then compared to the 99th percentile of the permuted samples to test whether the probability of the measured differences between different groups occurring by chance was smaller than 0.01.

### fMRI data analysis

fMRI data were processed using BrainVoyager QX (Brain Innovations, Maastricht, the Netherlands), MATLAB (MathWorks), SPM12^60^, the decoding toolbox^61^, and custom scripts. The anatomical volume in the retinotopic mapping session was transformed into the AC-PC space and then inflated using BrainVoyager QX. Functional volumes in all sessions were preprocessed, including 3D motion correction, linear trend removal, and high-pass filtering (cut-off frequency: 0.015 Hz) using BrainVoyager QX. The functional volumes were then aligned to the anatomical volume in the retinotopic mapping session and transformed into the AC-PC space.

The first 6 s of BOLD volumes were discarded to minimize transient magnetic saturation effects. A general linear model (GLM) procedure was used to define the ROIs. The ROIs in V1-V4 and LO were identified as a set of contiguous voxels that responded more strongly to the checkerboard stimulus than the blank screen (p<0.05, FDR corrected) in the localizer run.

With the fMRI data in the pre- and post-training tests, decoding analysis was performed to classify the activation patterns evoked by two orientations. A GLM procedure was first used to estimate beta values for individually responsive voxels in each stimulus block. The 100 most responsive voxels of each ROI were chosen to conduct the decoding analysis. The beta values of these voxels were then normalized in each run, resulting in 32 patterns (corresponded to 32 stimulus blocks) per test for each orientation. These patterns were used to train linear support vector machine (SVM) classifiers. A leave-one-run-out cross-validation procedure was used to calculate the decoding accuracy. Specifically, 28 patterns from 7 of 8 runs were chosen to train the classifier, and the remaining 4 patterns from the test run were used to estimate the decoding accuracy. The decoding improvement was calculated as (*Acc*_*post-training test*_ − *Acc*_*pre-training test*_)/*Acc*_*pre-training test*_ × 100%, where *Acc* stands for decoding accuracy.

FMRI data in the first training session of the dyadic training group was first analyzed using a univariate method to examine whether there was any difference in overall BOLD activity between the single and dyadic training runs. For each participant, a GLM was applied to the preprocessed functional imaging data. The GLM contained 2 regressors for the training types (single or dyadic) and 9 nuisance regressors accounting for run effects. Linear contrasts of estimated voxel-wise parameters were computed at the individual level and then taken to group-level *t* tests.

A searchlight analysis was then implemented to uncover any potential difference in neural activation pattern between the single and dyadic training runs. FMRI data during the single and dyadic training were firstly normalized to the template brain. Then we performed a volume-based searchlight analysis by sliding a 4-voxel-radius spherical linear SVM classifier voxel-by-voxel over the whole brain. The activity patterns within each searchlight were used to decode the training type by using a leave-one-run-out cross-validation procedure. To identify brain regions where the decoding accuracy was significantly higher than chance level, we performed second-level analyses by using voxel-wise one-sample t-tests on smoothed accuracy maps with a 3-D Gaussian kernel of 4 mm FWHM. This generated a voxel-wise whole-brain t-map reflecting the statistical significance of the decoding accuracy. Significance of the test was determined at uncorrected *p* < 1×10^−5^, cluster size *k* > 50.

Furthermore, we conducted PPI analysis to uncover how the functional connectivity between a given ROI (seed region) and a target region is modulated by a psychological variable. A 3D-Gaussian kernel of 6 mm FWHM was firstly used to smooth the EPI images. The seed brain regions were determined based on the searchlight analysis above. We used as a seed the average BOLD time series from a 3-voxel-radius spherical ROI in the bilateral IPL, lMFG, and ldlPFC, centered at the peak coordinates from the searchlight analysis mentioned above. Next, we constructed the interaction regressor of the PPI analysis (i.e., the regressor of main interest) by combining the seed ROI signals with the training type (dyadic = 1, single = −1) variable. The interaction regressor, together with the physiological (the BOLD time series of the seed region) and psychological (the training type variable) regressors entered into the first-level PPI design matrix. We further included 9 nuisance regressors accounting for run effects. The resulting first-level interaction regressor from each participant was then submitted to a second-level t test to establish the group-level connectivity maps. At last, we identified the overlapped regions of the group-level connectivity maps that derived from the seeds of ldlPFC, rIPL, and lIPL, respectively (uncorrected *p*<0.01).

## References

1. Bandura, A. & Walters, R. H. Social learning theory. vol. 1 (Englewood cliffs Prentice Hall, 1977).

2. Hill, G. W. Group versus individual performance: Are N+ 1 heads better than one? Psychological bulletin 91, 517 (1982).

3. Johnson, R. T. & Johnson, D. W. Active learning: Cooperation in the classroom. The annual report of educational psychology in Japan 47, 29–30 (2008).

4. Burke, C. J., Tobler, P. N., Baddeley, M. & Schultz, W. Neural mechanisms of observational learning. Proceedings of the National Academy of Sciences 107, 14431–14436 (2010).

5. Hammar Chiriac, E. Group work as an incentive for learning–students’ experiences of group work. Frontiers in psychology 5, 558 (2014).

6. Shteynberg, G., Hirsh, J. B., Bentley, R. A. & Garthoff, J. Shared worlds and shared minds: A theory of collective learning and a psychology of common knowledge. Psychological review 127, 918 (2020).

7. Ballard, I. C. et al. Dorsolateral prefrontal cortex drives mesolimbic dopaminergic regions to initiate motivated behavior. Journal of Neuroscience 31, 10340–10346 (2011).

8. Hayashi, T., Ko, J. H., Strafella, A. P. & Dagher, A. Dorsolateral prefrontal and orbitofrontal cortex interactions during self-control of cigarette craving. Proceedings of the National Academy of Sciences 110, 4422–4427 (2013).

9. Zhang, L. & Gläscher, J. A brain network supporting social influences in human decision-making. Science Advances 6, eabb4159 (2020).

10. Sasaki, Y., Nanez, J. E. & Watanabe, T. Advances in visual perceptual learning and plasticity. Nat Rev Neurosci 11, 53–60 (2010).

11. Watanabe, T. & Sasaki, Y. Perceptual Learning: Toward a Comprehensive Theory. Annu. Rev. Psychol. 66, 197–221 (2015).

12. Dosher, B. & Lu, Z.-L. Visual perceptual learning and models. Annual Review of Vision Science 3, 343–363 (2017).

13. Polat, U., Ma-Naim, T., Belkin, M. & Sagi, D. Improving vision in adult amblyopia by perceptual learning. Proceedings of the National Academy of Sciences 101, 6692–6697 (2004).

14. Maniglia, M. et al. Perceptual learning leads to long lasting visual improvement in patients with central vision loss. Restorative neurology and neuroscience 34, 697–720 (2016).

15. Barbot, A. et al. Spared perilesional V1 activity underlies training-induced recovery of luminance detection sensitivity in cortically-blind patients. Nature communications 12, 1–18 (2021).

16. Schoups, A. Practising orientation identification improves orientation coding in V1 neurons. Nature 412, 5 (2001).

17. Zohary, E., Celebrini, S., Britten, K. H. & Newsome, W. T. Neuronal plasticity that underlies improvement in perceptual performance. Science 263, 1289–1292 (1994).

18. Hua, T. et al. Perceptual Learning Improves Contrast Sensitivity of V1 Neurons in Cats. Current Biology 20, 887–894 (2010).

19. Adab, H. Z. & Vogels, R. Practicing coarse orientation discrimination improves orientation signals in macaque cortical area v4. Current biology 21, 1661–1666 (2011).

20. Jehee, J. F. M., Ling, S., Swisher, J. D., van Bergen, R. S. & Tong, F. Perceptual Learning Selectively Refines Orientation Representations in Early Visual Cortex. Journal of Neuroscience 32, 16747–16753 (2012).

21. Bi, T., Chen, J., Zhou, T., He, Y. & Fang, F. Function and Structure of Human Left Fusiform Cortex Are Closely Associated with Perceptual Learning of Faces. Current Biology 24, 222–227 (2014).

22. Lim, S. et al. Inferring learning rules from distributions of firing rates in cortical neurons. Nature neuroscience 18, 1804–1810 (2015).

23. Yu, Q., Zhang, P., Qiu, J. & Fang, F. Perceptual learning of contrast detection in the human lateral geniculate nucleus. Current Biology 26, 3176–3182 (2016).

24. Grill-Spector, K., Henson, R. & Martin, A. Repetition and the brain: neural models of stimulus-specific effects. Trends in Cognitive Sciences 10, 14–23 (2006).

25. Su, J., Chen, C., He, D. & Fang, F. Effects of face view discrimination learning on N170 latency and amplitude. Vision research 61, 125–131 (2012).

26. Chen, N., Lu, J., Shao, H., Weng, X. & Fang, F. Neural mechanisms of motion perceptual learning in noise. Human brain mapping 38, 6029–6042 (2017).

27. Dosher, B. & Lu, Z.-L. Perceptual learning reflects external noise filtering and internal noise reduction through channel reweighting. Proceedings of the National Academy of Sciences 95, 13988–13993 (1998).

28. Bejjanki, V. R., Beck, J. M., Lu, Z.-L. & Pouget, A. Perceptual learning as improved probabilistic inference in early sensory areas. Nature neuroscience 14, 642 (2011).

29. Gu, Y. et al. Perceptual learning reduces interneuronal correlations in macaque visual cortex. Neuron 71, 750–761 (2011).

30. Zivari Adab, H. & Vogels, R. Practicing Coarse Orientation Discrimination Improves Orientation Signals in Macaque Cortical Area V4. Current Biology 21, 1661–1666 (2011).

31. Law, C.-T. & Gold, J. I. Neural correlates of perceptual learning in a sensory-motor, but not a sensory, cortical area. Nat Neurosci 11, 505–513 (2008).

32. Law, C.-T. & Gold, J. I. Reinforcement learning can account for associative and perceptual learning on a visual-decision task. Nat Neurosci 12, 655–663 (2009).

33. Kahnt, T., Grueschow, M., Speck, O. & Haynes, J.-D. Perceptual learning and decision-making in human medial frontal cortex. Neuron 70, 549–559 (2011).

34. Dosher, B., Jeter, P., Liu, J. & Lu, Z.-L. An integrated reweighting theory of perceptual learning. Proceedings of the National Academy of Sciences 110, 13678–13683 (2013).

35. Rokem, A. & Silver, M. A. Cholinergic Enhancement Augments Magnitude and Specificity of Visual Perceptual Learning in Healthy Humans. Current Biology 20, 1723–1728 (2010).

36. Sereno, M. I. et al. Borders of multiple visual areas in humans revealed by functional magnetic resonance imaging. Science 268, 889–893 (1995).

37. Engel, S. A., Glover, G. H. & Wandell, B. A. Retinotopic organization in human visual cortex and the spatial precision of functional MRI. Cerebral cortex (New York, NY: 1991) 7, 181–192 (1997).

38. Wandell, B. A., Dumoulin, S. O. & Brewer, A. A. Visual Field Maps in Human Cortex. Neuron 56, 366–383 (2007).

39. Ni, Y. & Li, J. Neural mechanisms of social learning and decision-making. Science China Life Sciences 1–14 (2020).

40. Fliessbach, K. et al. Social comparison affects reward-related brain activity in the human ventral striatum. science 318, 1305–1308 (2007).

41. Gibbons, F. X. & Buunk, B. P. Individual differences in social comparison: development of a scale of social comparison orientation. Journal of personality and social psychology 76, 129 (1999).

42. Collins, R. L. For better or worse: The impact of upward social comparison on self-evaluations. Psychological bulletin 119, 51 (1996).

43. Hennig, J. A. et al. Learning is shaped by abrupt changes in neural engagement. Nature Neuroscience 1–10 (2021).

44. Luo, Y., Eickhoff, S. B., Hetu, S. & Feng, C. Social comparison in the brain: A coordinate-based meta-analysis of functional brain imaging studies on the downward and upward comparisons. Human brain mapping 39, 440–458 (2018).

45. Fehr, E. & Camerer, C. F. Social neuroeconomics: the neural circuitry of social preferences. Trends in cognitive sciences 11, 419–427 (2007).

46. Bhatt, M. A., Lohrenz, T., Camerer, C. F. & Montague, P. R. Neural signatures of strategic types in a two-person bargaining game. Proceedings of the National Academy of Sciences 107, 19720–19725 (2010).

47. McDonald, K. R., Pearson, J. M. & Huettel, S. A. Dorsolateral and dorsomedial prefrontal cortex track distinct properties of dynamic social behavior. Social cognitive and affective neuroscience 15, 383–393 (2020).

48. Mah, L., Arnold, M. C. & Grafman, J. Impairment of social perception associated with lesions of the prefrontal cortex. American Journal of Psychiatry 161, 1247–1255 (2004).

49. Chiao, J. Y. et al. Neural representations of social status hierarchy in human inferior parietal cortex. Neuropsychologia 47, 354–363 (2009).

50. Kedia, G., Lindner, M., Mussweiler, T., Ihssen, N. & Linden, D. E. Brain networks of social comparison. Neuroreport 24, 259–264 (2013).

51. Uddin, L. Q., Molnar-Szakacs, I., Zaidel, E. & Iacoboni, M. rTMS to the right inferior parietal lobule disrupts self–other discrimination. Soc Cogn Affect Neurosci 1, 65–71 (2006).

52. Seitz, A. R., Kim, D. & Watanabe, T. Rewards evoke learning of unconsciously processed visual stimuli in adult humans. Neuron 61, 700–707 (2009).

53. Wang, Z., Kim, D., Pedroncelli, G., Sasaki, Y. & Watanabe, T. Reward Evokes Visual Perceptual Learning Following Reinforcement Learning Rules. bioRxiv 760017 (2019).

54. Arsenault, J. T., Nelissen, K., Jarraya, B. & Vanduffel, W. Dopaminergic reward signals selectively decrease fMRI activity in primate visual cortex. Neuron 77, 1174–1186 (2013).

55. Katz, B., Shah, P. & Meyer, D. E. How to play 20 questions with nature and lose: Reflections on 100 years of brain-training research. Proceedings of the National Academy of Sciences 115, 9897–9904 (2018).

56. Levi, D. M. & Li, R. W. Perceptual learning as a potential treatment for amblyopia: a mini-review. Vision research 49, 2535–2549 (2009).

57. Yang, X.-Y., He, Q. & Fang, F. Transcranial direct current stimulation over the visual cortex facilitates awake consolidation of visual perceptual learning. Brain Stimulation: Basic, Translational, and Clinical Research in Neuromodulation 15, 380–382 (2022).

58. He, Q., Yang, X.-Y., Gong, B., Bi, K. & Fang, F. Boosting visual perceptual learning by transcranial alternating current stimulation over the visual cortex at alpha frequency. Brain Stimulation 15, 546–553 (2022).

59. Brainard, D. H. The Psychophysics Toolbox. Spatial vision 10, 433–436 (1997).

60. Penny, W., Friston, K., Ashburner, J. & Kiebel, S. Statistical Parametric Mapping: The Analysis of Functional Brain Images. (2007). doi:10.1016/B978-0-12-372560-8.X5000-1.

61. Hebart, M. N., Görgen, K. & Haynes, J.-D. The decoding toolbox (TDT): A versatile software package for multivariate analyses of functional imaging data. Frontiers in neuroinformatics 8, (2015).

62. Bex, P. J. & Makous, W. Spatial frequency, phase, and the contrast of natural images. JOSA A 19, 1096–1106 (2002).

63. Pelli, D. G. & Bex, P. Measuring contrast sensitivity. Vision research 90, 10–14 (2013).

64. Foster, D. H. & Bischof, W. F. Thresholds from psychometric functions: superiority of bootstrap to incremental and probit variance estimators. Psychological Bulletin 109, 152 (1991).

